# TEDlm: domain-centric protein language models with optional structural pre-training

**DOI:** 10.64898/2026.07.09.737428

**Authors:** Tiejun Wei, Shaun M. Kandathil, Daniel W. A. Buchan, David T. Jones

**Author notes:** Corresponding author: David T. Jones.

## Abstract

Conventional protein language models are pretrained on full-length sequences that interleave multiple domains with linkers and disordered regions, diluting fold-specific signals. Our approach pretrains masked language models on structurally-defined domain segments from The Encyclopedia of Domains. TEDlm learns from domain sequences alone with a standard MLM objective, while its variant TEDlm3D adds a Cα distance-guided contact loss that supervises the attention maps. On CATH S40 remote-homology detection (<40% identity), the domain-centric pretraining has a bigger effect than model scale: at the final layer, a 650M-parameter TEDlm achieves an AUROC1 of 0.28 compared to 0.22 for ESM2 3B, whereas TEDlm3D reaches 0.50, approaching the structure-based search tool Foldseek (0.53) from sequence alone at inference. Attention-map and categorical Jacobian probes show that the contact signal is encoded in the model representations themselves, not only in a trained output head. TEDlm variants also substantially improve zero-shot Molecular Function prediction over ESM2, while matching it on various biophysical property tasks, indicating that signals are largely domain-intrinsic. Together, these results position domain-centric pretraining as a route to compact, structurally informed protein language models.

## Introduction

Proteins underpin essentially all biological processes yet inferring their three-dimensional structures and molecular functions directly from amino acid sequences remains a fundamental challenge despite recent breakthroughs in deep learning–based structure prediction^1,2^. Genome and metagenome sequencing projects have produced billions of sequences, creating urgent demand for scalable methods to detect homology and infer structure and function *in silico*^3^. Alignment-based tools such as BLAST, HMMER, and HHblits have long been used for this task^4–6^. Recent algorithms like MMseqs2 significantly improved the speed and sensitivity of sequence searches, with better sensitivity than PSI-BLAST while being hundreds of times faster^7^. However, pure alignment-based methods struggle when sequence identity approaches the so-called twilight zone^8,9^, leading to the development of machine learning methods that can detect subtle sequence signals of structural or functional similarity. To date, accurate homology detection remains a crucial first step for transferring structural and functional annotations between proteins.

The advent of transformer architectures^10^, which revolutionized natural language processing through contextual self-attention and large-scale pretraining^11^, has similarly reshaped bioinformatics. Deep transformer-based protein language models (pLMs) have achieved remarkable success in capturing evolutionary and structural patterns from self-supervised pretraining on large-scale sequence datasets^12–15^. The success of sequence-only pLMs may be attributed to their ability to learn the evolutionary information embedded in protein sequences, commonly reflected as conserved regions that have accumulated over millions of years. The Evolutionary Scale Models (ESM) family of pLMs demonstrated these evolutionary patterns can be captured by pretraining on unaligned sequences^16,17^. Learning rich protein representations was thought to be achievable only through relatively expensive computation of multiple sequence alignments (MSAs)^18^.

Remarkably, large pLM embeddings contain signals predictive of secondary structure, residue–residue contacts, and protein function, despite being trained only to model sequences^15^. A landmark example is ESM2, a masked language model whose embeddings, learned from unaligned sequences alone, carry enough structural information that ESMFold predicts atomic-level protein structure from a single sequence^16^. Recently, embedding-based methods such as PLMSearch have demonstrated that even frozen pLMs, e.g. the modestly sized ESM-1b model, can rival alignment-based methods like MMseqs2 for homology detection with a simple cosine similarity ranking^7,15,19^. These findings underscore the evolutionary signal captured in pretrained sequence embeddings alone, without any structural supervision.

The rapid expansion of structural data in the AlphaFold Protein Structure Database (AFDB)^1,3^ has enabled another approach: bimodal sequence-structure pLMs. ProstT5, a structure-aware model trained on top of ProtT5, incorporates explicit structural tokens alongside sequence tokens^12,13^. SaProt and ProSST take a related route, the Foldseek 3Di alphabet in SaProt and a learned quantized alphabet in ProSST; both require an explicit structure at inference^20,21^. Another recent approach, TM-Vec/DeepBLAST, explicitly incorporates structural information by training on TM-align scores as an auxiliary loss, producing embeddings that correlate more directly with structural similarity^22^. Although the combination of sequence and structure tokens significantly boosts homology detection performance, it also relies on ground-truth structures during training^21^. Similarly, Foldseek leverages a predefined vocabulary of 20 local residue environments called 3Di tokens, which enables ultra-fast structural similarity search while retaining high sensitivity in homology detection^23^.

Independently of these bimodal approaches, Rao et al. demonstrated that transformer attention maps trained with an unsupervised masked language modelling objective learn residue–residue contacts directly, achieving state-of-the-art unsupervised contact prediction from a single forward pass^24^. More recently, Zhang et al. showed through a categorical Jacobian analysis that pLMs such as ESM-2 store statistics of coevolving residue motifs analogously to Markov Random Field models, though the structural signal captured remains limited by having been trained on full-length sequences^25^. These findings suggest that the self-attention mechanism in pLMs is a natural architecture for encoding residue-level geometric relationships, and that explicitly guiding this mechanism with structural supervision during pretraining could amplify the contact signal beyond what unsupervised objectives achieve alone.

Although the growing availability of protein structures enables explicit incorporation of structural information, significant potential remains in improving sequence-based pretraining alone. Many proteins are composed of two or more domains, which are compact, independently folded functional units of protein sequence and structure. Conventional pLMs are typically trained on full-length sequences, which may contain several concatenated domains, often in different orders^13,16,20^. This could obscure the learning of domain-specific patterns, as the model is trained on sequences of longer average length, which contain unstructured/linker regions. Recent breakthroughs in deep learning-based protein domain segmentation, such as Merizo^26^, ChainSaw^27^, and UniDoc^28^, have made it possible to effectively segment and classify protein domains in data sets as large as the AFDB. A comprehensive dataset of domains in the AFDB is now available, known as The Encyclopedia of Domains (TED)^29^. It identified more than 365 million domains, with ∼100M more domains than were detectable by sequence-based methods. By segmenting full-length proteins into domain-level sequences, models can be trained on structurally coherent, functionally minimal regions, potentially capturing more implicit structural signals from sequence alone, as compared to training on full-length protein sequences.

In this work, we introduce TEDlm and TEDlm3D, transformer-based protein language models that leverage segmented domain sequences during pretraining to introduce fold-relevant inductive bias. TEDlm adopts a BERT-like masked language model objective and learns to predict masked amino acids within domain sequences, capturing motifs and long-range dependencies that define specific folds and superfamilies. TEDlm3D extends this framework by incorporating a Cα distance-guided contact prediction auxiliary loss, exploiting the observation that transformer attention maps naturally encode residue–residue proximity. By jointly optimising sequence prediction and contact geometry on domain structures derived from the AFDB, TEDlm3D is designed to explicitly encode the structural relationships that TEDlm captures only implicitly. The key hypothesis is that learning from domain-level sequences, optionally augmented with structural supervision, enables the model to disentangle structure-relevant patterns from extraneous sequence noise, yielding representations that generalise across structural and functional prediction tasks.

Finally, we examine how structural information is distributed across transformer layers. On homology detection, intermediate-layer embeddings substantially outperform final-layer ones across every model we test; the gap is large enough that even TEDlm3D 160M, from sequence input alone, surpasses Foldseek at an intermediate layer. This mirrors observations in natural language processing (NLP), where intermediate layers encode richer representations than the final layer and consistently outperform final-layer embeddings on downstream tasks^30^. We interpret this as the final layers of a masked language model specialising for the token-level prediction objective, compressing the more abstract, fold-discriminative features carried in intermediate layers. A systematic layer-wise analysis across all models is given in Supplementary Table 1.

## Results

### Domain pretraining and structural auxiliary loss improves fold recognition

We assessed TEDlm and TEDlm3D on homology detection using the CATH S40 dataset, a canonical benchmark for evaluating a model’s ability to capture structural relationships at low sequence identity (<40%)^31^. Sensitivity was measured in AUROC1 (area under the ROC curve up to the first false positive), which quantifies how reliably true remote homologs are ranked ahead of non-homologs. We compared against alignment-based methods (MMseqs2, Foldseek) and state-of-the-art pLMs (ESM2, ProtT5, ProstT5)^7,12,13,16,23^. All pLMs were evaluated using exhaustive all-vs-all cosine similarity ranking; alignment-based tools used exhaustive search mode.

Restricting pretraining to domain segments improved fold discrimination on its own. Using final-layer embeddings, TEDlm 650M outperformed both ESM2 models, the alignment tool MMseqs2, and matching encoder-only ProtT5 3B. A 650M domain model therefore surpasses a full-length sequence model more than four times its size, hinting that on the homology detection task, the pretraining data outweighs model scale. TEDlm3D, which combines the same domain-based pretraining with the Cα distance-guided contact objective, shows further improvement in accuracy from sequence input alone, approaching the structure-based tool Foldseek. Notably, the TEDlm3D 160M variant exceeded every sequence-only baseline, while remaining competitive with ProstT5 3B, which uses structural tokens during training.

The structural signal was spread across the network rather than concentrated at the output (Figure 1B). Every model performs substantially better with intermediate layer embeddings compared to their final layer embeddings on this task, indicating that the final layer understates the structural information and is optimised for its MLM sequence loss objective. ESM2 models have the biggest gap with its final-layer performance falling far below its intermediate-layer peak. TEDlm3D models (650M and 1.3B) lose much less and interestingly even their final-layer embeddings exceed the best layer of either ESM2 model. As an upper reference, TEDlm 1.3B w/CATH, which adds a CATH topology classification loss, reaches AUROC1 = 0.733; because this loss directly shapes the embedding geometry that AUROC1 measures, we report it as weakly supervised metric learning rather than a zero-shot result. Per-layer values are listed in Supplementary Table 1.

**Figure 1.**
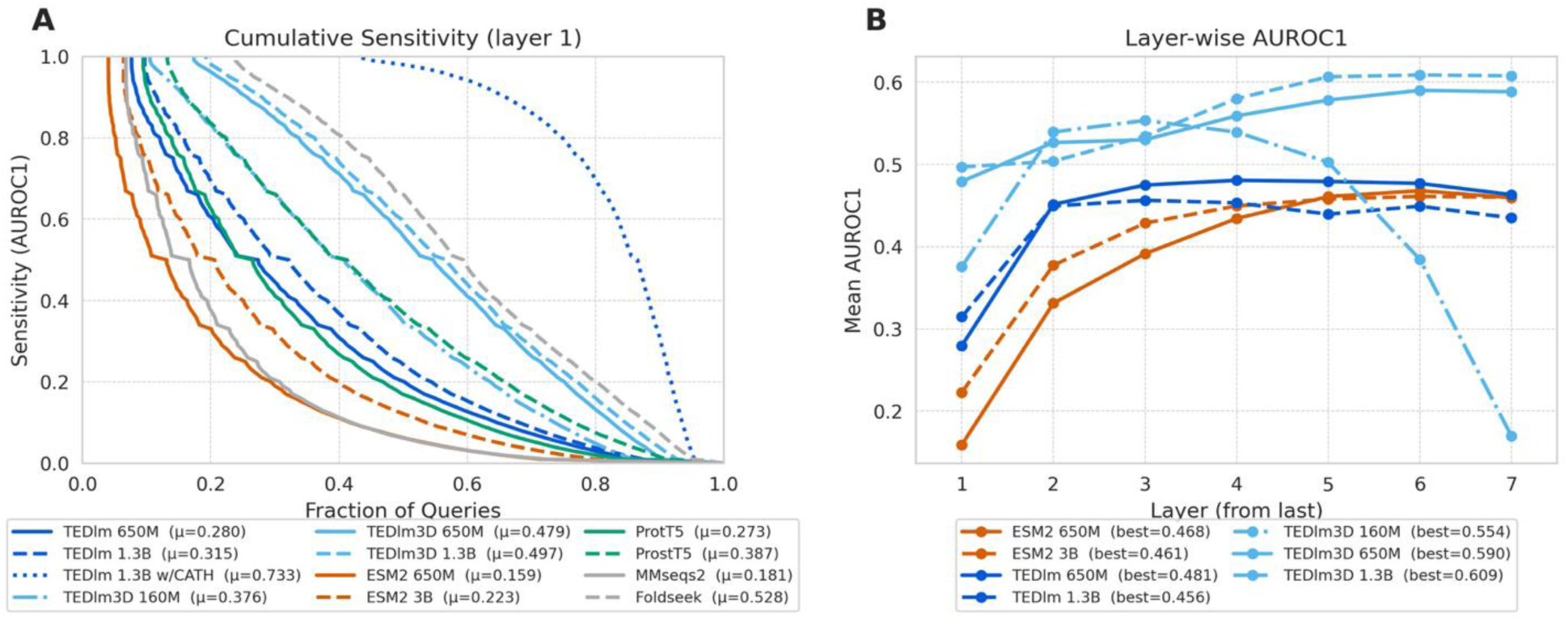
Homology detection performance on the CATH S40 dataset. (A) Cumulative sensitivity distributions using final-layer embeddings. True positives are matches within the same CATH superfamily; false positives are matches between different folds. Mean AUROC1 (μ) is shown in the legend. (B) Layer-wise AUROC1 across the last seven transformer layers (layer 1 = final layer). Best AUROC1 achieved at any layer is indicated in the legend.

### TEDlm3D contact head captures residue-level structural information

We tested whether the Cα objective leaves a residue-level contact signal in the learned representations rather than only in a trained output layer, using two probes on 1,327 CATH S40 v4.4.0 domains, one per topology (Methods). We first probed the attention maps, which reflect internal information mixing rather than the output logits (Figure 2A). TEDlm3D and TEDlm 650M share the same architecture and the MLM objective and differ in the Cα loss only. The best long-range head of TEDlm3D 650M reached precision at sequence length (P@L) = 0.347 against 0.205 for TEDlm 650M, and 0.304 against 0.244 at short and medium range, with the effect concentrated in a few heads rather than the attention apparatus (mean long-range P@L ≈ 0.04 across all 672 heads; Methods). The Cα loss thus reshapes how TEDlm3D mixes residue information internally rather than adding a separate readout.

**Figure 2.**
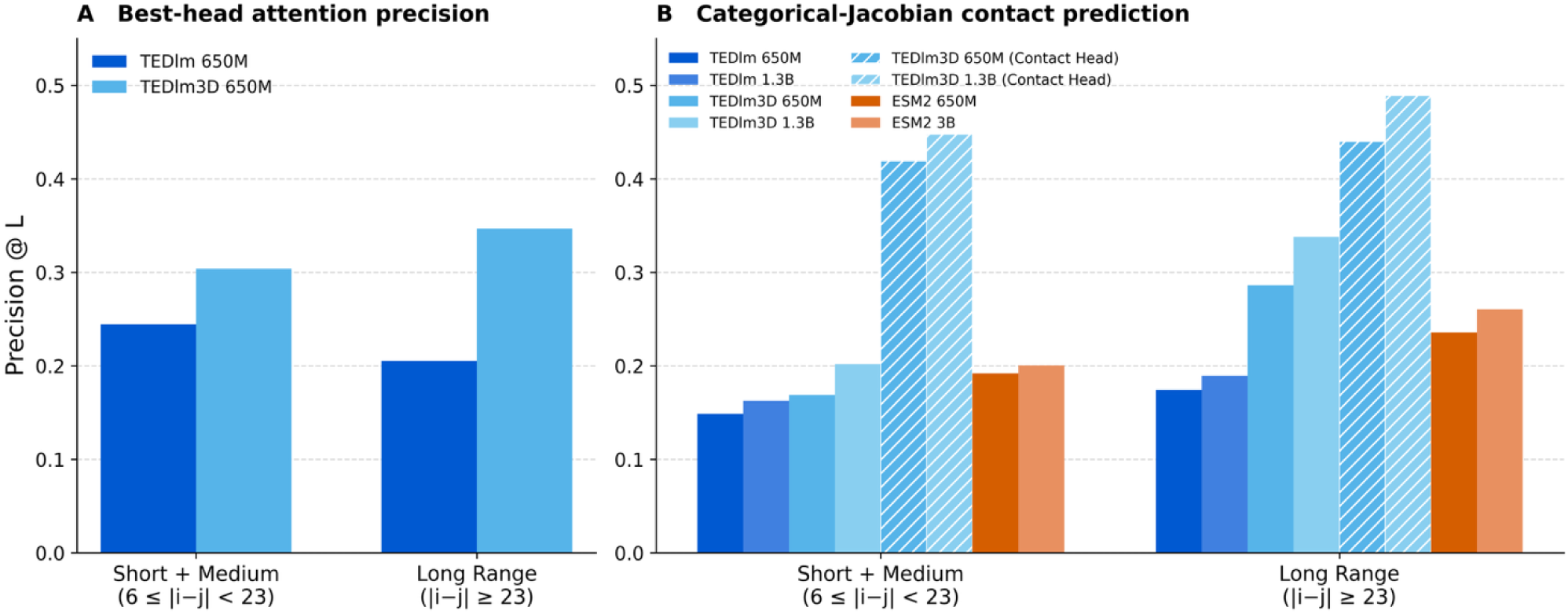
TEDlm3D captures long-range structure better than TEDlm under two independent probes. Precision@L on 1,327 CATH topology representatives, for residue pairs split into short+medium (6 ≤ |i−j| < 23) and long-range (|i−j| ≥ 23) bins. (A) Best-head attention precision for the TEDlm 650M / TEDlm3D 650M pair. (B) Categorical-Jacobian contact prediction across all models (hatched bars: TEDlm3D supervised contact head). Both panels showed TEDlm3D performs better than its original in terms of contact prediction, while the advantage widens at long range. **Unsupervised contact prediction precision** (Ovchinnikov Jacobian method) and TEDlm3D contact head precision across P@L/5, P@L/2, and P@L thresholds. Left: medium- and long-range contacts (6 < |i−j| <23); right: long-range only (|i−j| ≥ 23). Solid bars denote unsupervised Jacobian-derived predictions; hatched bars denote TEDlm3D contact head predictions. Contacts defined as Cα distance < 8 Å, with APC correction applied.

The same pattern held end-to-end under the categorical Jacobian of each model’s logits, which scores coevolutionary coupling without any trained contact layer and applies uniformly across architectures (Figure 2B)^25^. For long-range pairs (|i−j| ≥ 23), TEDlm3D 1.3B reached P@L/5 = 0.552, above ESM2 3B (0.438) and its MLM-only counterpart TEDlm 1.3B (0.348), with the same ordering at smaller sizes. The contact signal recoverable from a single forward pass is therefore stronger in TEDlm3D than in an equally sized full-length model or in TEDlm.

Within TEDlm3D models, a dedicated contact head, trained on Cα distance maps during pretraining, reads this signal out at high precision in one forward pass (hatched bars, Figure 2B). For medium- and long-range pairs it reached P@L/5 = 0.745 (650M) and 0.789 (1.3B), against about 0.45 for the best unsupervised Jacobian baseline (ESM2 3B); for long-range pairs it reached 0.638 and 0.698 at P@L/5, and 0.440 and 0.489 at the stricter P@L. Note the head is also an order of magnitude faster than the unsupervised Jacobian approach, which requires a forward pass per residue substitution.

### Training on domain sequences yields better performance on MF Prediction

We evaluated zero-shot GO term prediction by KNN label transfer on two benchmarks: the time-cutoff CAFA3 test^32^, whose knowledge base predates the CAFA3 targets and excludes homologs (Table 2), and an up-to-date Swiss-Prot test that stratifies queries by maximum sequence identity to the knowledge base (Figure 3; Methods). Molecular function (MF) was the namespace where domain pretraining helped consistently. The gap between models on the CAFA3 targets was large: on MF namespace TEDlm and TEDlm3D variants exceeded ESM2, with TEDlm 1.3B at wFmax = 0.336 and TEDlm 650M at 0.323 against ESM2 3B (0.301) and ESM2 650M (0.270). The Swiss-Prot test confirmed the trend: across the identity-capped bins from 0.1 to 0.8, every TEDlm and TEDlm3D variant exceeded both ESM2 models, and at a 0.3 cap, TEDlm 650M and 1.3B reached 0.488 and 0.486 against ESM2 3B (0.474) and ESM2 650M (0.458), a lead near 0.01 to 0.015 that closed to parity with ESM2 3B only when all neighbours were admitted. TEDlm and TEDlm3D tracked each other on MF, with TEDlm ahead at most thresholds, so the gain reflects the domain training data rather than the contact objective. Notably, most enzymes realize their function within an individual domain, while a minority of active sites form at the interface between two or more domains^33^. The molecular-function gains we observe are therefore consistent with domain-level pretraining capturing function-relevant signal.

**Figure 3.**
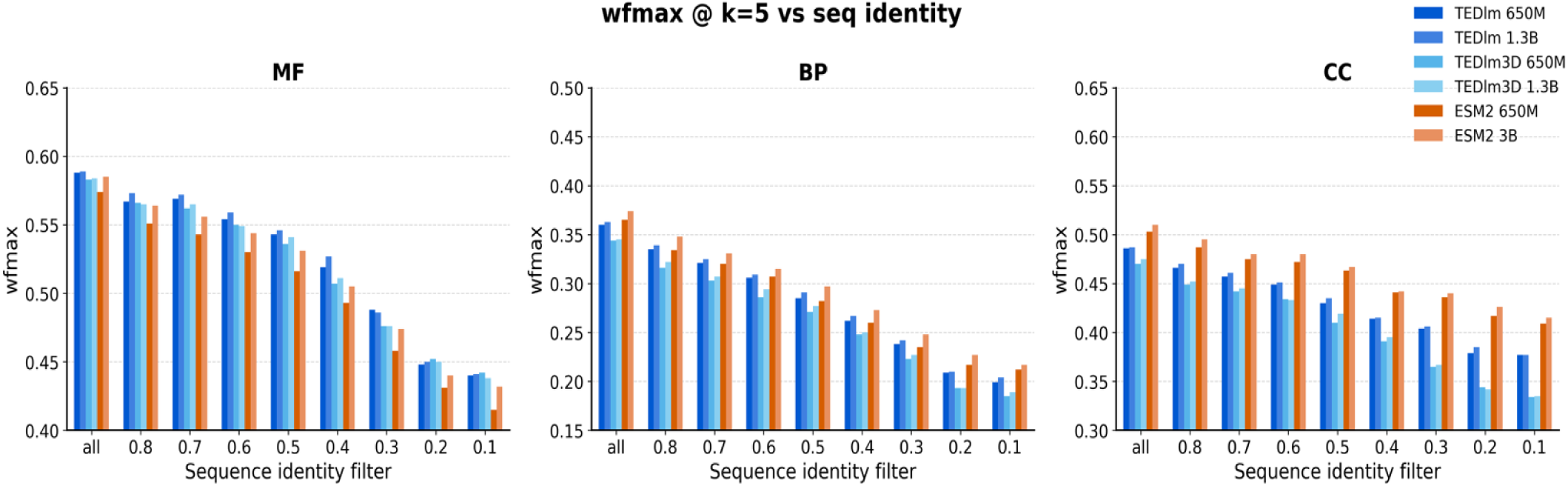
Zero-shot KNN-based GO term prediction. (wFmax at K = 5) on the customized UniProtKB-SwissProt benchmark across three ontology namespaces (MF, BP, CC). X-axis indicates the maximum allowed query–neighbor sequence identity, with additional E-value < 10^−3^ training mask applied at each threshold.

**Table 1:**
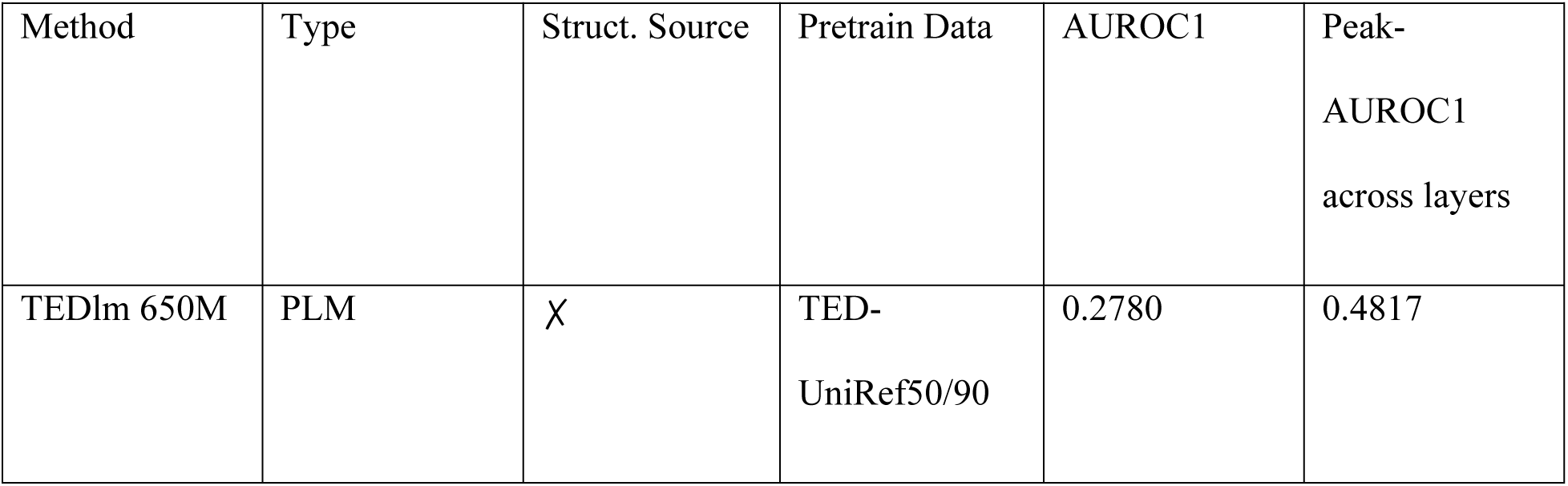
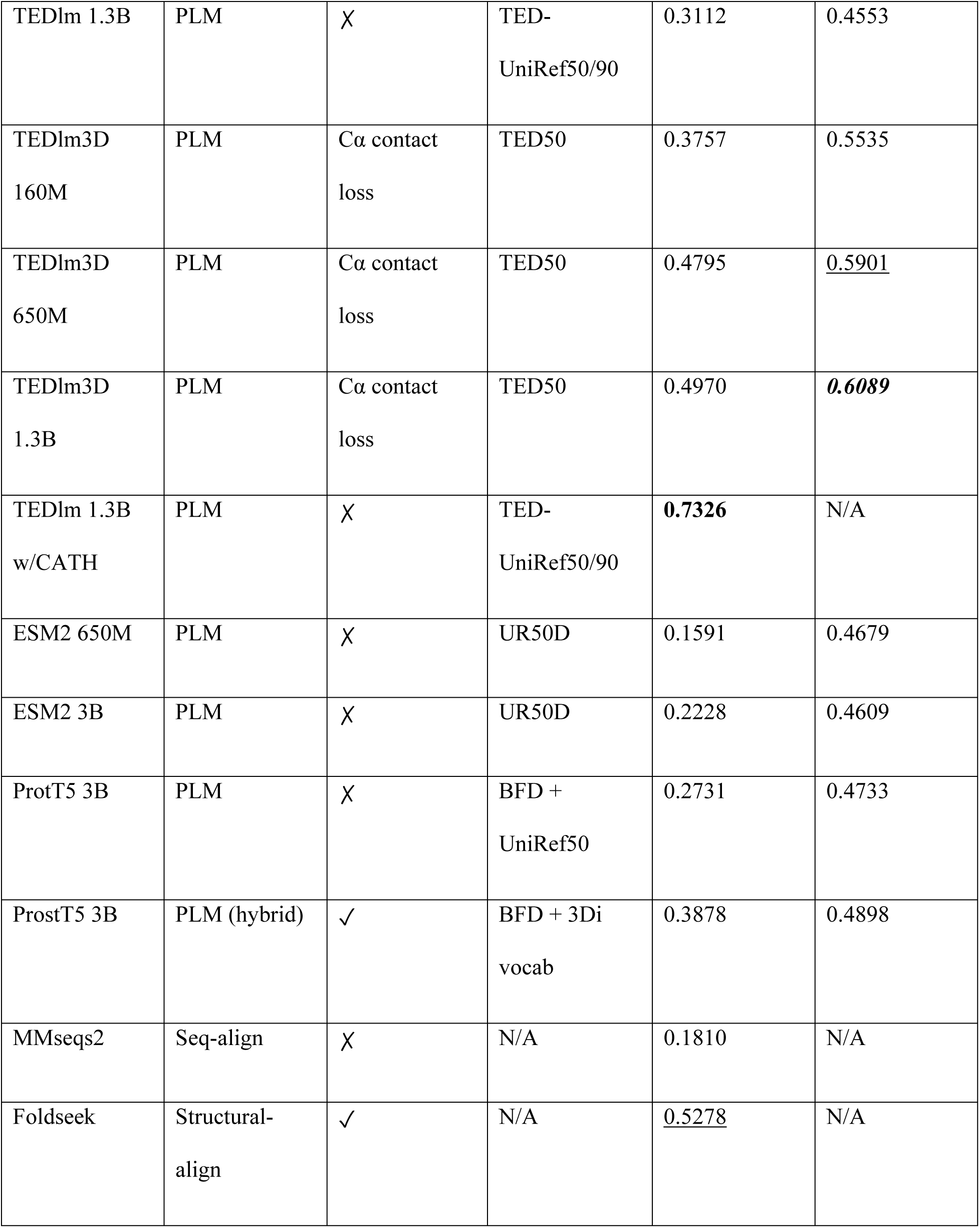
Homology detection (AUROC1) performance on the CATH S40 dataset. The canonical AUROC1 uses final-layer embeddings; peak-AUROC1 reports the peak across the last seven transformer layers. Best overall in bold; second-best underlined.

**Table 2:** Zero-shot GO term prediction performance (weighted Fmax) on the CAFA3 benchmark with k=5. Homologous sequences excluded by E-value < 10. Best per namespace in bold. Error bar represents half-width of a 95% bootstrap confidence interval (10,000 protein-level resamples).

| Ontology | MF wFmax | BP wFmax | CC wFmax |
| --- | --- | --- | --- |
| TEDlm 650M | 0.323 ± 0.021 | 0.136 ± 0.010 | 0.287 ± 0.019 |
| TEDlm 1.3B | **0.336 ± 0.021** | 0.131 ± 0.009 | 0.270 ± 0.018 |
| TEDlm3D 650M | 0.308 ± 0.020 | **0.182 ± 0.012** | 0.260 ± 0.017 |
| TEDlm3D 1.3B | 0.304 ± 0.020 | 0.157 ± 0.008 | 0.270 ± 0.018 |
| ESM2 650M | 0.270 ± 0.019 | 0.145 ± 0.007 | 0.294 ± 0.019 |
| ESM2 3B | 0.301 ± 0.021 | 0.138 ± 0.007 | **0.296 ± 0.018** |

In biological process (BP) the two benchmarks disagree, and we do not claim a domain advantage in either. On CC, ESM2 models remain leading position in the CAFA3 test (Table 2), on the up-to-date Swiss-Prot test, ESM2 held the advantage at every identity threshold (Figure 3), with ESM2 3B highest and the TEDlm3D variants lowest; at a 0.3 cap ESM2 3B reached BP wFmax 0.248 and CC 0.440, above TEDlm 1.3B (0.242 and 0.406). This fits the expectation that biological process and subcellular localisation are more likely to depend on multi-domain context, signal peptides and protein interactions that domain-level pretraining does not see.

### Domain pretraining achieves property prediction across biophysical tasks

To assess whether the representational advantages of domain-centric pretraining extend beyond homology detection, we evaluated all models on five biophysical property prediction tasks using a KNN label transfer protocol with sequence-identity-stratified evaluation (S. Figure. 6; see Methods). Tasks included enzyme catalytic efficiency (Kcat, Spearman ρ)^34^, metal-ion binding (AUROC)^35^, and thermostability prediction on human, cross-species, and nanobody datasets (Spearman ρ)^36–38^. For each task, an MMseqs2-based search was first used to restrict the dataset to single-domain proteins^7^. An additional training mask was then applied to exclude neighbours above a specified sequence-identity threshold, creating a series of increasingly stringent evaluation bins that probe whether embeddings capture transferable signals beyond high sequence similarity.

Across tasks, TEDlm and TEDlm3D achieved performance comparable to ESM2. TEDlm variants showed modest advantages under stringent filtering for enzyme catalytic efficiency, with Kcat at a 60% identity cap TEDlm 650M (0.45) outperforming ESM2 650M (0.36); and metal-ion binding (20% sequence identity cap), TEDlm 1.3B (0.74) v.s. ESM2 3B (0.66), where the relevant biophysical signals are defined at the domain level by active-site motifs and coordination geometries. TEDlm3D performed comparably to TEDlm on most tasks, but weaker on thermostability prediction, and the clearest case where ESM2 holds advantage was cross-species thermostability (TEDlm 650M ρ = 0.49 vs 0.54 for ESM2 650M), likely because organism-level thermal adaptations extend beyond individual domain properties. Taken together, properties governing catalysis and metal coordination appears largely intrinsic to individual domain sequences and can be captured without full-protein context^34–36^.

## Discussion

In this work, we introduced TEDlm and TEDlm3D, protein language models pretrained on structurally defined domain segments from The Encyclopedia of Domains (TED)^29^. Between TEDlm and ESM2, we have demonstrated the training dataset effect: restricting training to domain-level sequences alone produces representations that outperform models up to 4-fold larger on a stringent homology detection task. TEDlm 650M outperforms ESM2 3B on homology detection despite having fewer than a quarter of the parameters, and TEDlm3D, which adds a Cα distance-guided contact prediction objective, approaches the performance of Foldseek (AUROC1 = 0.497 vs 0.528) while requiring only sequence input at inference^7,16,23^. The layer-wise analysis showed that the weakest layer of TEDlm3D 650M and 1.3B still outperforms ESM2’s peak performance, so domain pretraining with structural supervision retains fold-relevant information up to the final layer rather than concentrating it in intermediate layers. The TEDlm 1.3B w/CATH variant, which incorporates CATH topology-level classification as an auxiliary loss, pushes this further (AUROC1 = 0.733), illustrating that domain-centric pretraining provides a strong foundation upon which even modest structural supervision yields disproportionate gains, a design principle shared by TEDlm3D through its complementary contact-level geometric loss.

The contact prediction benchmark provides direct mechanistic evidence for what TEDlm3D’s structural auxiliary loss encodes. The unsupervised categorical Jacobian approach shows that TEDlm3D representations contain richer coevolutionary signals than either TEDlm or ESM2^25^, and a dedicated contact head reads this out at high precision in a single forward pass. A common assumption in protein language modelling is that representations are dominated by short-range correlations. This is well motivated: in masked token prediction, much of the recoverable information is contained within local sequence context, such as motifs associated with secondary structure. Our results suggest that this can be addressed with auxiliary losses derived from structure. Although the contact prediction head shows strong performance across both short- and long-range interactions, as we might expect, we also find that the learned embeddings themselves encode substantially more long-range information than those of baseline models such as ESM, as demonstrated by the ability of the amino acid projections to better reconstruct long-range contacts. Crucially, the Cα objective supplies no structural input at inference; TEDlm3D still reads sequence alone, so it acts purely as a training-time constraint rather than an added information channel. Comparing TEDlm3D against TEDlm, the most closely matched pair available (Figure 2A), TEDlm3D’s attention maps recover long-range contacts more accurately (best head P@L 0.347 vs 0.205 at 650M), placing this gain in the learned representation itself rather than in a readout head trained on top of it. This suggests that the auxiliary contact loss directly influences the geometry of the representation space, encouraging the whole model to better incorporate long-range structural dependencies. Taken together, these findings suggest that incorporating structurally motivated auxiliary objectives can mitigate the intrinsic locality bias of masked language modelling and promote the learning of more globally informative representations. On the locally defined tasks, catalytic efficiency and metal coordination, TEDlm and TEDlm3D match or slightly exceed ESM2, which supports our interpretation that properties such as catalytic efficiency, metal coordination are better represented by single-domain trained TEDlm models. However, thermal stability prediction is the exception and seems to favour full-length models. The functional annotation benchmarks reveal both the strengths and the principled limits of domain-level representations. On the CAFA3 benchmark, all TEDlm and TEDlm3D variants substantially outperform ESM2 in predicting Molecular Function (wFmax up to 0.336 vs 0.301), consistent with the expectation that many MF annotations, (e.g. catalytic activities and binding specificities), are attributable to individual domains. The advantage was specific to MF. On the up-to-date Swiss-Prot benchmark, ESM2 led in both Biological Process and Cellular Component prediction across all identity thresholds, with TEDlm3D lowest, the two benchmarks diverging because CAFA3 removes close homologs and uses a dated knowledge base that rewards remote-homology signal.

On the other hand, this reflects an intrinsic granularity mismatch: GO annotations are curated at the gene-product level, while TEDlm is pretrained on isolated domain segments with no exposure to multi-domain architectures or protein complex contexts that often determine subcellular localisation. Domain-level embeddings may therefore require explicit aggregation mechanisms to recover protein-level CC assignments, as domain-informed annotation resources such as InterPro2GO have demonstrated^39^. Several limitations should be acknowledged. The domain boundaries inherited from TED, though derived from high-quality AlphaFold models and consensus segmentation tools^26–28^, may include boundary inaccuracies that propagate into pretraining. For TEDlm, the training data undergoes redundancy reduction at the protein level (UniRef50/90)^40^ before domain segmentation, meaning the actual redundancy structure of the domain-level training set has not been directly controlled. We addressed this by curating TED50/90 from the 365M complete TED dataset and trained TEDlm3D upon this. However, another limitation could arise from the fact that pretraining data is drawn from AFDB version 4^3^, inherently biasing toward high-confidence predictions and under-representing rare folds and intrinsically disordered regions. TEDlm does not incorporate multiple sequence alignments, foregoing the explicit evolutionary depth that has proven valuable in methods such as AlphaFold2^1,2^. Finally, the performance gains from intermediate embeddings, documented in the extended data, suggest that final transformer layers in pLMs are partially optimised for token-level prediction at the expense of global structural representations^30^, a phenomenon warranting further investigation. Future work should explore domain-level deduplication strategies, hierarchical representations bridging domain and protein levels for functional annotation, and integration of TEDlm3D’s domain-centric approach with multi-domain and complex-level context.

## Methods

### Data preparation

Using the 2021_04 release of UniProt, protein sequences were obtained by cross-referencing the UniProt identifiers from UniRef50 representative sequences (49,874,565 entries) and UniRef90 (138,515,945 entries) against version 4 of the AFDB (214,683,829 entries)^3,40^, yielding a customised set of 41,859,106 UniRef50 entries and 108,502,280 UniRef90 entries which could be exactly matched against AFDB structure models. Domain boundary annotations were sourced from version 5 TED release^29^. We then cross-referenced the custom UniRef50/90 sets against the consensus TED domain boundaries, retaining 36,458,420 entries with matching domain annotations. Each matched protein was then truncated into its constituent domains according to TED consensus boundaries, producing a curated set of compact, independently folded sequence segments for downstream transformer pretraining, which we call the TED- UniRef50 and TED-UniRef90 sets. 0.1% of the whole dataset was held out as a validation set, while the remainder was used as the training set. As a result, TED-UniRef50 has 60,064,241 entries in the training set and 60,125 in the validation set, whereas TED-UniRef90 has 177,642,528 and 177,821 entries, respectively.

We then constructed TED90 and TED50 directly from TED365, taken from The Encyclopedia of Domains. TED365 was first clustered with MMseqs2 at 90% sequence identity and 60% coverage, yielding 190,626,362 representative sequences (TED90) that constitute a redundancy-reduced pretraining dataset^7^. The TED90 representatives were then re-clustered at 50% identity (60% coverage) to produce TED50, with a total of 84,811,276 entries. As the cluster result is highly imbalanced, with a few super-hubs containing up to 21,749 members and the majority containing many fewer, we adopted a weighted training scheme whereby the learning rate is weighted by the inverse size of the cluster to address the pretraining dataset imbalance.

### Homology detection benchmark

We evaluated the effectiveness of TEDlm in capturing structural and evolutionary signals using a customized homology detection benchmark based on the CATH S40 dataset^31^. This benchmark is designed to assess a model’s ability to identify homology relationships between structurally related proteins with low sequence identity (<40%), in which traditional sequence alignment-based methods often underperform. For context, we benchmarked our model against several SOTA protein language models and other tools:

- the deep learning-based ESM2 models, namely esm2_t33_650M_UR50D, and esm2_t36_3B_UR50D, referred to as ESM2 650M and ESM2 3B respectively;
- ProtTrans, also known as ProtT5, with the 3B denoising objective-tuned checkpoint prot_t5_xl_uniref50, referred to as ProtT5 3B;
- ProstT5, an encoder-decoder style model trained bidirectionally with sequence and 3Di tokens upon the ProtT5-XL-U50 checkpoint, in its latest checkpoint, referred to as ProstT5 3B;
- Foldseek; and
- MMseqs2, a commonly used sequence alignment-based tool.

Note that all deep learning-based models are evaluated using a strict all-vs-all (exhaustive) similarity ranking test. The alignment-based tools, i.e. Foldseek and MMseqs2, have prefilters enabled but running in exhaustive search mode. This is to ensure all tools are performing at their best capability. The pLM models are ranked using average-pooled embedding cosine similarity, while bit score and qTM score were used for MMseqs2 and Foldseek, respectively. The ProtT5 and ProstT5 models use an encoder-only approach, thus effectively using ∼ 1.5B parameters. All pLM baselines are evaluated from sequence input, with Foldseek included as a structure-based upper reference; structure-token pLMs such as SaProt and ProSST require per-residue structural tokens at inference to fully leverage their difference to ESM2 models and were not trained for sequence-only inference mode, so they were not benchmarked. The selected methods serve as informative baselines of sequence-based embedding search, structure-sequence embedding search and alignment-based search in zero-shot style.

### Enzyme Catalytic efficiency, thermostability, and metal binding prediction

For the zero-shot enzyme catalytic efficiency, metal-ion binding, and thermostability benchmarks, we used the same CAFA3 style evaluation protocol described below, in which protein embeddings are used for KNN label transfer from the training set, i.e. the knowledge base or the “universe”, to test set. For enzyme catalytic efficiency, we used the Enzyme Catalytic Efficiency dataset hosted at Hugging Face (https://huggingface.co/datasets/biomap-research/enzyme_catalytic_efficiency), consisting of 16,838 entries. These measurements were originally curated from multiple enzyme-kinetics repositories by Li *et al.* (DLKcat) and subsequently adopted as a benchmark in xTrimoPGLM^34,41^. Spearman’s correlation coefficient between predicted values and experimental K_cat_ was used as the main metric for this task. Similarly, we used the metal-binding train/test set, where a binary classification task was performed via KNN, and AUROC was used as the main metric^35^. For thermostability, we curated task-specific splits from melting-temperature resources, including gene-level melting-point aggregation derived from the Meltome Atlas thermal proteome profiling study^36^. We cross-referenced both the human and cross-species meltome datasets against the TED365 domain database. Using MMseqs2, we performed sequence similarity searches with the easy-search module to identify hits with 80% coverage. For enzymatic efficiency prediction, metal binding and thermostability prediction tasks, sequences were searched against the TED365 fasta file with MMseqs2 suite, easy-search mode with coverage of 80% and sequence similarity of 80%, to form a single-domain subset of data. A random split with train/validation/test ratio of 8:1:1 was created. Nanobody thermostability data were adopted from NbThermo and NanoMelt^37,38,42^. Detailed number of dataset entries can be found in S. Table 2.

To reduce leakage from near-identical sequences, including homologues and point-mutation variants, we applied an MMseqs2-based training mask during KNN label propagation using a series of maximum allowed sequence-identity thresholds^7^. Any neighbour above the specified sequence-identity threshold was excluded from query label voting before the cosine-similarity search. All results were generated with k=5, last layer embedding in the heat map S. Figure 6.

### Contact prediction from single-sequence Jacobians

We assessed the capacity of each model to encode residue–residue contact information using the categorical Jacobian approach of Zhang et al^25^. For a protein of length L, we constructed a full (L × 20 × L × 20) Jacobian tensor by substituting each position i with all 20 canonical amino acids and recording the change in predicted logits at every other position j relative to the wild-type sequence. The resulting tensor was centred by iteratively subtracting the mean along each of its four axes, collapsed to an (L × L) coupling matrix via the Frobenius norm over the two amino-acid dimensions, symmetrized, and corrected for phylogenetic and positional biases with the average product correction (APC)^43^. For TEDlm3D models, we additionally evaluated a supervised contact head: a single forward pass through the model captured the final hidden representations h ∈ ℝ^ (L × d), which were projected through a learned linear layer. The (L × L) contact-probability matrix was then obtained by element-wise sigmoid followed by symmetrization. APC was not applied to the contact-head output, as the supervised head already accounts for positional biases.

The benchmark was evaluated on 1,327 CATH S40 (v4.4.0) domains selected as one representative per topology (T-level). Domains were restricted to 50–300 residues containing only canonical amino acids. Ground-truth binary contact maps were derived from Cβ atomic coordinates (Cα for glycine) at an 8 Å distance threshold. We report precision at L/5, L/2, and L top-scoring predictions, as well as the area under the receiver operating characteristic curve (AUROC), stratified by sequence separation into short/medium range (6 ≤ |i − j| < 23) and long range (|i − j| ≥ 23).

### Per-head attention precision

TEDlm and TEDlm3D use Flash Attention, which does not expose attention weights. For each sequence we registered a forward pre-hook on every attention layer’s inner kernel to capture the post-rotary query and key tensors and recomputed the attention matrix explicitly as scaled-dot-product-attention (SDPA) via softmax(QKᵀ/√d) per head. Specifically, for both 650M models: 28 layers × 24 heads = 672 heads. To ensure the attention matrix remained the same as Flash Attention, we recomputed weights and the maximum absolute error < 5×10⁻³ across both models. Each head’s L×L attention matrix was symmetrised (A + Aᵀ) and APC-corrected, then scored by precision against the gold labels. Only the best performing head was selected, i.e. the head with the highest mean long-range P@L over the 1,327 CATH S40 T-level representatives. For TEDlm: layer 27/head 5; and for TEDlm3D: layer 25/head 15, the Figure 2A reports that head’s precision in both short+medium and long-range bins. As a negative control, the mean long-range P@L across all 672 heads is around 0.04 for both models, differing drastically to the heads with high mean long-range P@L (0.347 for TEDlm3D 650M and 0.205 for TEDlm 650M).

### GO prediction benchmark

We evaluated the model’s ability to generalize functional relationships using a modified, stringent, zero-shot style of the CAFA3 benchmark^32^. Functional annotation transfer is fundamentally related to homology detection, as both depend on identifying sequence relationships that reflect shared structural or evolutionary ancestry. Unlike standard KNN-style evaluations, our custom setting deliberately removes any homologous proteins from the training knowledge base, creating a far more stringent test of generalization. This design probes whether representations learned through the domain-centric pretraining could capture meaningful functional similarity when sequence similarity is low. To construct the training set, or the knowledge base used during the KNN, we used the intersection between Swiss-Prot and the UniProt-GOA 2017_01, prior to the date preceding the CAFA3 release (2017_01)^40,44^. During the KNN-based label propagation, any sequence with identical UniProt ID was excluded. MMseqs2 easy search with a sensitivity of 30 and query/target coverage of 60% was conducted, and any query with E-value smaller than 10 was also excluded^7^. Embeddings for both the knowledge base and target sequences were computed using the same approach as in the homology detection benchmark, and functional prediction was performed using cosine-similarity–weighted label propagation within the Molecular Function (MF), Biological Process (BP), and Cellular Component (CC) namespaces.

To further examine how embedding similarity relates to functional similarity, we randomly sampled 100,000 non-homologous protein pairs from the Swiss-Prot and UniProt-GOA 2017_01 set, excluding any pairs with E-values smaller than 10. Each sequence was embedded with the aforementioned models, and mean pooling was applied across residues to obtain a single vector representation per protein.

Pairwise cosine similarity scores were computed and normalized to the [0,1] range, with the embedding distance defined as D, where s° is the cosine similarity min-max scaled to [0,1], as shown in equation (2). To assess functional similarity, we computed the Jaccard Index between the GO term sets of each pair across the namespaces, using GO annotations derived from the 2017_01 Gene Ontology Annotation (GOA) release^44,45^.

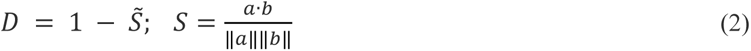

The functional similarity between two proteins A and B was quantified using the Jaccard Index J(A, B), defined as:

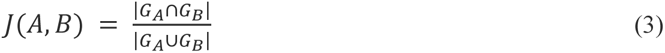

where G_A_ and G_B_ denote the sets of GO terms annotated to proteins A and B, respectively^45^.

AUROC was used to quantify the model’s ability to distinguish functionally related from unrelated domain pairs. A pair was considered functionally similar when its GO Jaccard index exceeded 0.5, corresponding to a positive label.

For the UniProtKB-SwissProt benchmark set, we start by collecting sequences with experimental-only labels, the allowed annotation evidence codes are EXP, IDA, IPI, IMP, IGI, IEP, HTP, HDA, HMP, HGI, HEP. The resulting set is then cross-referenced with the latest GOA, yielding ∼79k sequences. To avoid data leakage, we chose the cluster-then-random-assign approach. We started by clustering sequences based on different thresholds of sequence identity in [0.1, …, 0.9], then randomly assigned the member sequences and their representatives into the train/test sets, with a soft constraint on label balancing and achieving roughly the ratio of 8:2 for the train and test sets^46^. The label balancing is done by randomly assigning the train-test set 100 times with a fixed random seed for reproducibility. A term is considered as valid when it is present in both train and test set, and the random assignment with the most valid label pairs are selected for use as the benchmark. This is done across the sequence identity range. To prevent leakage between near-identical sequences lying near cluster boundaries, we ran an all-against-all MMseqs2 search over the training set and generated a training mask, which effectively masked any pair with E-value below 0.001 during the KNN label transfer. A CAFA6 information accretion file, as provided on the CAFA6 official website, is used for the weighted F_max_ (wF_max_) and weighted S_min_ (wS_min_) calculation using the cafa-evaluator tool^47^.

## Supporting information

supplementary matrials

## Contributions

D.T.J. conceived the study, the domain-centric pretraining approach and carried out model training. D.T.J designed and implemented the TEDlm and TEDlm3D architectures. T.W. assembled the domain-level pretraining dataset from TED, curated and performed the downstream evaluation and analysis. T.W. prepared the figures and wrote the original draft. S.M.K. and D.W.A.B. contributed to method development, benchmarking, and evaluation design. D.T.J. supervised the project and acquired funding. All authors reviewed and edited the manuscript and approved the final version.

## Data Availability

The pre-trained weights for all TEDlm and TEDlm3D variants are deposited on Zenodo https://zenodo.org/records/20814105.

## Code Availability

Code for model inference and benchmark is available at https://github.com/psipred/tedlm under the Apache-2.0 licence. A versioned snapshot of the repository corresponding to this publication is archived on Zenodo https://zenodo.org/records/20814105.

## Acknowledgements

We thank Dr Owain Kenway for valuable discussions and comments. This work was supported by the Biotechnology and Biological Sciences Research Council (BBSRC) BB/T019409/1. We gratefully acknowledge the computational resources and support provided by UCL Central Cluster Myriad, Advanced Research Computing (ARC), and the use of the UCL Department of Computer Science high-performance computing cluster for model development and training. This work also used the Isambard-AI National AI Research Resource (AIRR). Isambard-AI is operated by the University of Bristol and funded by the UK Government’s Department for Science, Innovation and Technology (DSIT) via UK Research and Innovation (UKRI) and the Science and Technology Facilities Council (STFC) 0251-9628-2113-1.

## Ethics declarations

### Competing interests

No competing interests statement is required.

## Supplementary Information

Supplementary Material provided separately.

## Notes

https://zenodo.org/records/20814105

### Competing Interest Statement

The authors have declared no competing interest.

