## supplementary matrials for "TEDlm: domain-centric protein language models with optional structural pre-training"

### TEDlm Supplementary Materials

#### Neural Network Architecture

TEDlm models use transformer encoder architectures trained with a masked language modelling objective. The 650M-parameter model represents each input token as a 1,536-dimensional vector and processes it through 28 transformer layers with multi-head self-attention. To improve training efficiency, it uses Rotary positional embeddings with FlashAttention-2[1,2], together with padding/unpadding strategies that avoid unnecessary attention calculations on padding tokens. Each transformer block applies pre-layer normalization, followed by attention and a SwiGLU feed-forward module, before a final normalization and linear prediction head produce logits for masked token prediction[3].

The larger TEDlm 1.3B model follows the same overall design but increases the model width to 2,048 dimensions, uses 36 transformer layers, and has 32 attention heads. The TEDlm 1.3B w/CATH variant adds an auxiliary loss that trains the model to predict topology-level (T-level) CATH labels alongside the masked language modelling task.

The TEDlm3D models keeps the TEDlm backbone unchanged and adds a structure-aware second head. A linear projection maps each residue embedding to a structure vector, whose pairwise inner products serve as logits predicting a Cα–Cα contact map. The head is supervised on contacts read from AlphaFold structures with a hard filter of pLDDT >= 0.7 (or 70 out of 100). We applied uncertainty weighting on this Cα auxiliary loss with the masked language model loss so the model balances sequence and geometry signals automatically during the pre-training[4]. For details see model training section below.

#### Model training

For training, we adopted a two-phase training procedure with all data points sampled with equal weights. As outlined in the network structure section, the model was trained with a masked language modelling objective, i.e. predicting the identity of amino acids that have been randomly masked out of protein sequences. The <eod> token was added at each domain sequence end.

$$L_{MLM}(\theta) = -\frac{1}{|M|}\sum_{i \in M} \log P(\theta)(x_i|x_M) \qquad \text{(S1)}$$

As shown in Equation S1, for a randomly generated mask *M* that includes 15% of positions *i* in the sequence x, the model was tasked with predicting the identity of the amino acids $x_i$ in the mask from the surrounding context $x_M$, excluding the masked positions. The model was trained with ~60M domain sequences in the TED-UniRef50 set for 13 epochs, then ~177M domain sequences with TED-UniRef90 set for 3 epochs until the validation loss plateaued. We adopted the AdamW optimizer with a learning rate of 0.003, as determined by torch lightning tuner module, weight decay is set to 0.001[5,6]. The batch size was set to 48 to maximize the GPU memory utilization, and gradients were clipped to 1.0 for better training stability. Training was carried out in fp16 mixed precision on an 8-way NVIDIA A6000 node for approximately 1 month, with the model seeing ~100B tokens over the course of training to be on par with ESM2 650M model training[7].

TEDlm3D has an additional loss term $L_{C\alpha}(\theta)$, fomulated as below:

$$L_{C\alpha}(\theta) = BCE(z_{i,j}, y_{i,j}) \qquad \text{(S2)}$$

$$L_{C\alpha}(\theta)' = \frac{\sum_{i,j} w_{i,j} L_{C\alpha}(\theta)}{\sum_{i,j} w_{i,j}} \qquad \text{(S3)}$$

$$L_{total} = \frac{L_{MLM}(\theta)}{2e^{s_{MLM}}} + \frac{L_{C\alpha}(\theta)}{2e^{s_{C\alpha}}} + \frac{s_{MLM} + s_{C\alpha}}{2} \qquad \text{(S4)}$$

Where the C$\alpha$ loss term is simply the predicted distogram $z_{i,j}$ with BCE loss evaluated regarding the ground truth distogram $y_{i,j}$, per equation S2[8]. This is weighted with a hard filter on pLDDT > 0.7 minimum on both residue in the i,j pair, then normalized, resulting in filtered C$\alpha$ loss, shown as in equation S3. The total loss for the TEDlm3D used during pretraining shown in Equation S4. which learns each task's relative weight from its estimated uncertainty, while the ½s terms prevent the model from trivially suppressing either task.

Regarding the training time and resources, TEDlm3D 160M model was trained with NVIDIA A6000 node for approximately two weeks, 650M model was trained with 8 AMD MI300X GPUs for approximately a week, while the TEDlm3D 1.3B model was trained with 16 GH200 GPUs on the UK's AI Research Resource (AIRR) for approximately 2 weeks.

**Loss-function ablation at matched data and scale (160M)**

The contact objective raises the masked-language-modelling cross-entropy above the BERT-only baseline (middle panel) while the contact loss converges smoothly (right panel), showing that the two objectives compete for shared-trunk capacity. The auxiliary signal therefore acts as a structural constraint at a small cost to token prediction. We briefly benchmarked the homology and various downstream property prediction performance with the two models, see result below.

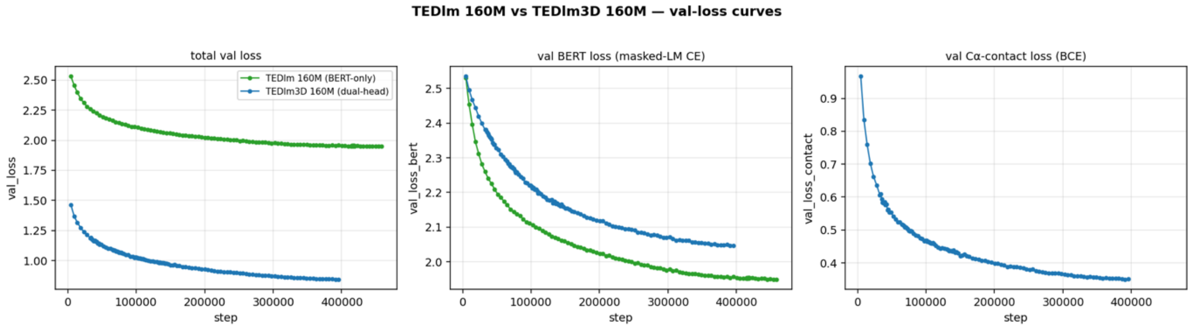


***S. Figure 1.** Validation loss for the 160M data-matched pair. TEDlm 160M (BERT-only, green line) and TEDlm3D 160M (dual-head, blue line) were trained on the identical TED50 corpus with the same architecture, optimizer and compute budget, differing only in the addition of the Cα contact auxiliary loss. Three panels showed: total validation loss, for TEDlm this is simply the MLM loss, for TEDlm3D this is the combined loss after the weighting (left panel); MLM loss (middle). Cα contact loss defined only for the dual-head model(right). Curves are computed on the held-out TED50 validation split.*

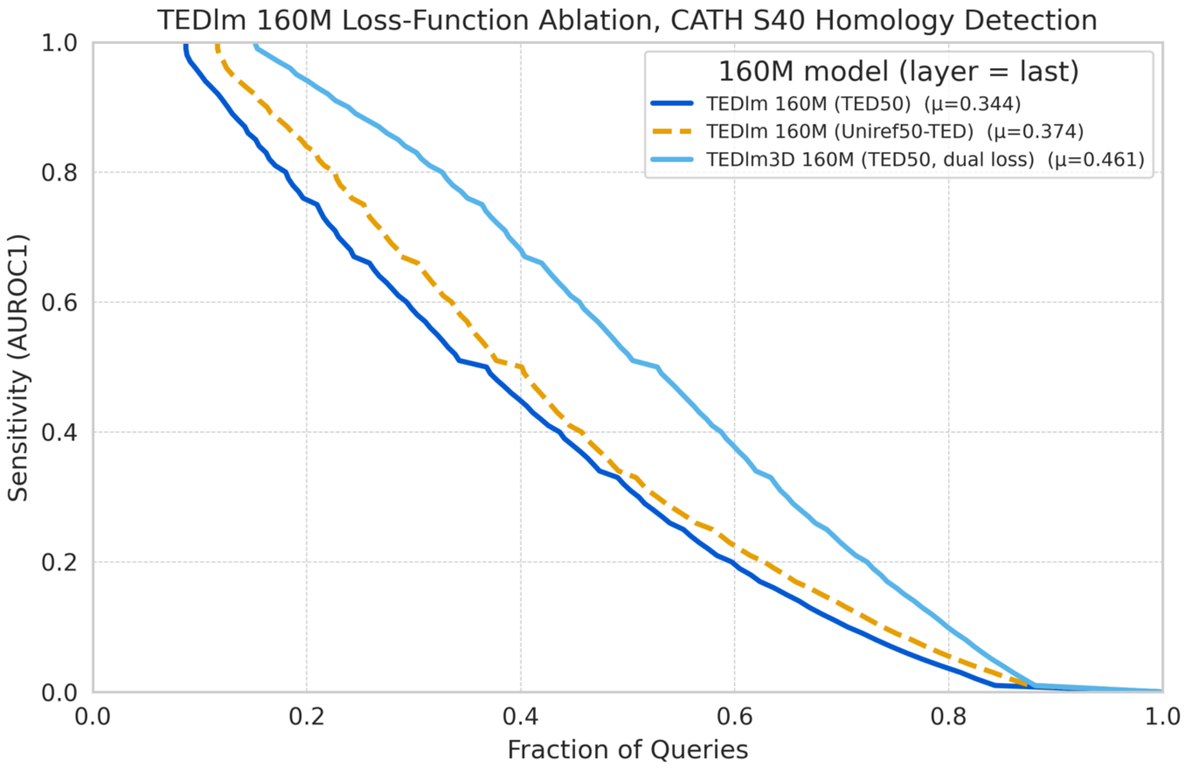


***S. Figure 2. Loss-function and corpus ablation on CATH S40 homology detection.*** *Cumulative sensitivity using final-layer embeddings (AUROC1 shown as μ in the legend) for three 160M models: TEDlm trained on TED50 (dark blue solid line, μ = 0.344), TEDlm trained on TED-UniRef50 (orange dashed line, μ = 0.374), and TEDlm3D trained on TED50 with the joint masked-language-modelling and Cα contact loss (light blue solid line, μ = 0.461). The TED50 trained TEDlm and TEDlm3D models are matched on corpus, architecture and compute, isolating the effect of the contact objective; the two TEDlm models share the objective and differ only in corpus. True positives are matches within the same CATH superfamily; false positives are matches between different folds (for details see CATH S40 benchmark, Methods).*

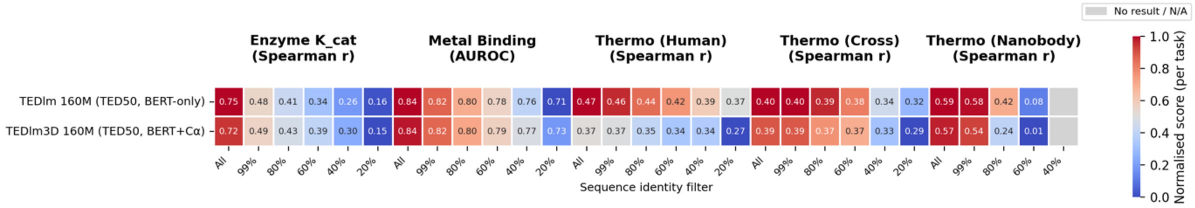


***S. Figure 3. Biophysical property prediction is preserved under the contact objective.*** *Zero-shot KNN label transfer (k = 5, final-layer embeddings) for TEDlm 160M (TED50, BERT-only) and TEDlm3D 160M (TED50, BERT + Cα) across five tasks. Cell values are Spearman ρ for enzyme $K_m$ efficiency (Kcat) and the human, cross-species and nanobody thermostability tasks, and AUROC for metal-ion binding; color is normalized per task. Grey cells indicate thresholds with too few retained pairs to score.*

**S. Table 1:** Homology detection performance on CATH S40 test set (AUROC1) with various intermediate layer embedding. KNN retrieval using embeddings computed from embeddings extracted from the last k transformer blocks (-1 to -8, where -1 denotes the final transformer block). Best AUROC1 per model is shown in **bold.** Here we can see the trend the best embedding for fold recognizing task depends on its relative position/depth within the model rather than an arbitrary layer index.

| Model | Layer-1 | Layer-2 | Layer-3 | Layer-4 | Layer-5 | Layer-6 | Layer-7 | Layer-8 |
|---|---|---|---|---|---|---|---|---|
| TEDlm 650M | 0.2782 | 0.4519 | 0.4761 | **0.4817** | 0.4799 | / | / | / |
| TEDlm 1.3B | 0.3112 | 0.4409 | 0.4547 | **0.4553** | 0.4406 | 0.4513 | / | / |
| TEDlm3D 160M | 0.3758 | 0.5396 | **0.5535** | 0.5394 | 0.5026 | 0.3846 | 0.1696 | / |
| TEDlm3D 650M | 0.4795 | 0.5267 | 0.5301 | 0.5591 | 0.5784 | **0.5902** | 0.5885 | / |
| TEDlm3D 1.3B | 0.4970 | 0.5039 | 0.5345 | 0.5800 | 0.6067 | **0.6089** | 0.6078 | / |
| ESM2 650M | 0.1590 | 0.3314 | 0.3915 | 0.4344 | 0.4609 | **0.4680** | 0.4596 | 0.4557 |
| ESM2 3B | 0.2227 | 0.3774 | 0.4286 | 0.4495 | 0.4578 | **0.4609** | 0.4600 | 0.4566 |
| ProtT5 (ProtTrans) | 0.2731 | 0.2373 | 0.3048 | 0.3819 | 0.4293 | 0.4541 | 0.4652 | **0.4733** |
| ProtsT5 | 0.3878 | 0.3359 | 0.3955 | 0.4355 | 0.4634 | 0.4790 | 0.4871 | **0.4898** |

#### CATH S40 homology overlap analysis against TED50/Uniref50

To quantify benchmark–training redundancy for the CATH S40 homology-detection task, we compared the 34,653 CATH non-redundant S40 domain sequences against the pre-training data set of TEDlm3D (TED50). CATH S40 queries were searched against TED50 with MMseqs2 with -s 5.7 -e 10 --max-seqs.

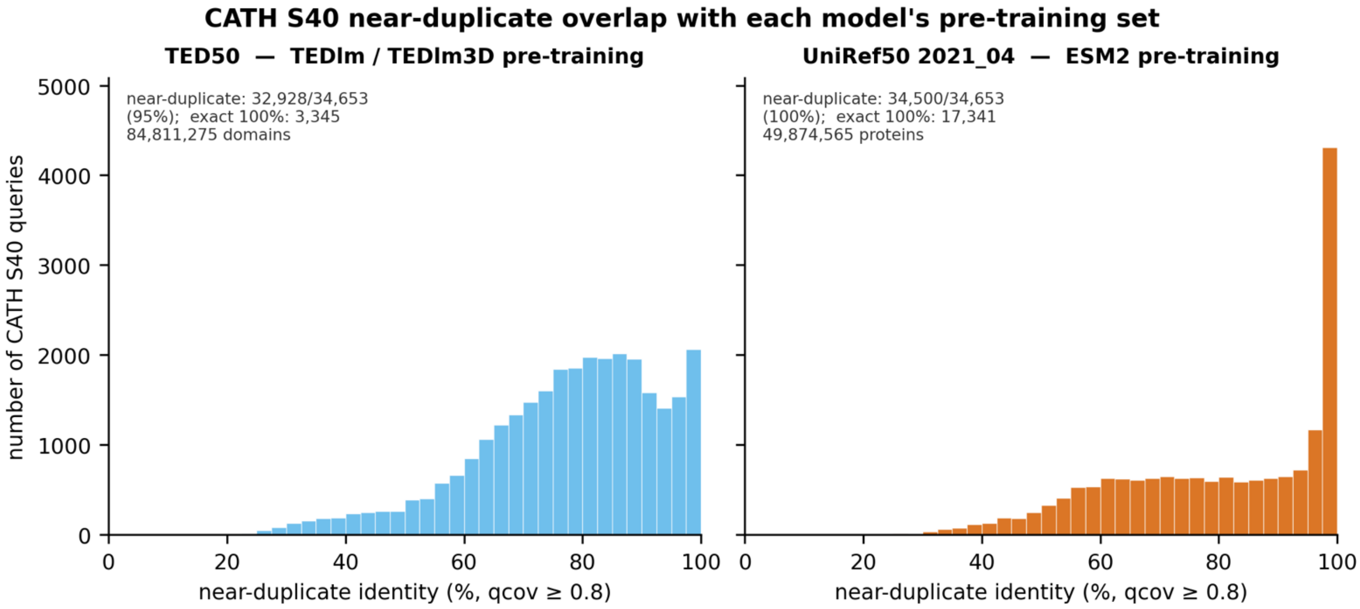


***S. Figure 4.*** *Distribution of the closest near-duplicate sequence identity between each CATH S40 query and the TED50/Uniref50. Where nearly half of CATH S40 entries are 100% match in the Uniref50 2021_04 set, which ESM2 training set UR50D was curated on.*

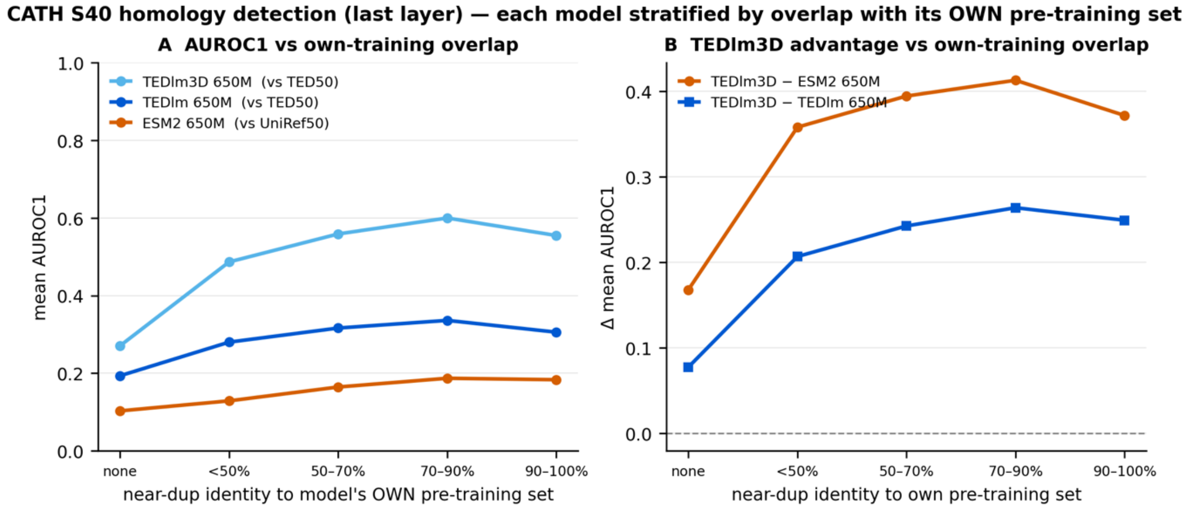


***S. Figure 5.*** CATH S40 homology-detection AUROC1 from last-layer embeddings, with queries stratified by the sequence identity of their closest TED50 near-duplicate.

We then stratified the AUROC1 performance of each CATH S40 query according to their closest homologue as identity bins. Not surprisingly the performance of TEDlm and TEDlm3D models shrink considerately when the test cases get harder (to the left side of the plot). However, we’d like to point out even in the none bin the ordering holds, TEDlm3D still outperforms other models, with AUROC1 of 0.25 over TEDlm 0.18 over ESM2 0.14. Indicating domain pretraining and the contact loss do generalise to folds model never saw before. Note that there are only 97 entries in the ‘none’ bin for ESM2, so the data point is noisy, but we included it to show the trend only.

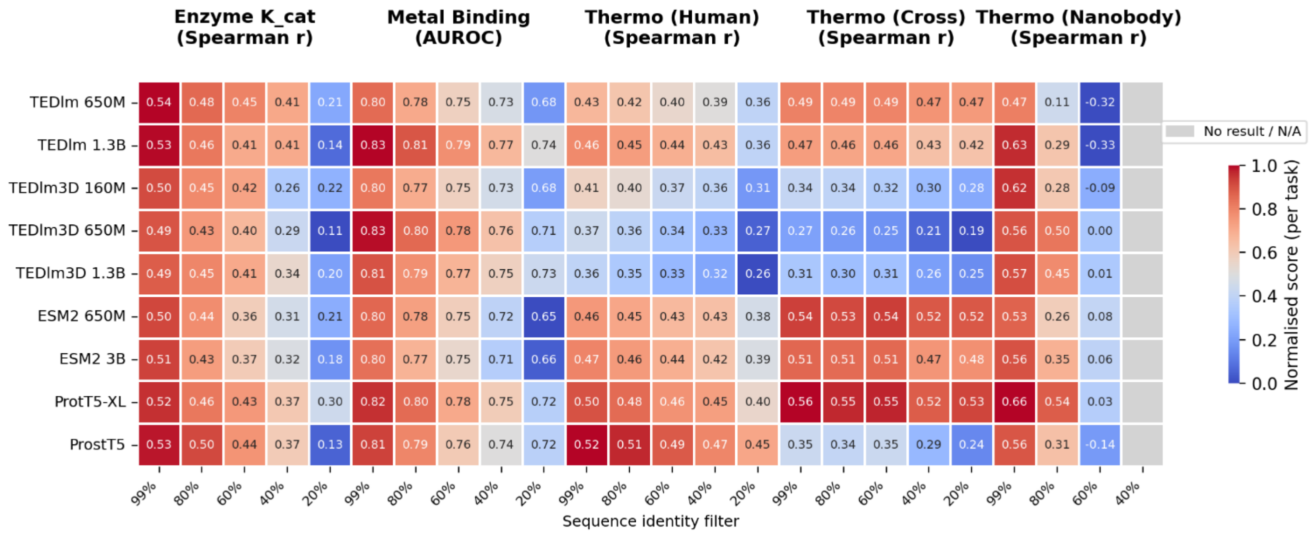


***S.Figure 6 Single-domain Zero-shot property prediction via KNN label transfer with sequence-identity filtering.*** *Heatmaps report performance across five biophysical tasks for TEDlm, TEDlm3D, ESM2, ProtT5-XL, and ProstT5 models. Columns indicate the maximum allowed query–neighbor sequence identity. Metrics: Spearman ρ for enzyme catalytic efficiency ($K_{cat}$), human thermostability, cross-species thermostability, and nanobody thermostability; AUROC for metal-ion binding. Color scale is normalized per task. For each model–task–threshold combination, we used k=5, last layer embedding.*

| Task | Canonical set (train/val/test) | Single-domain set (train/test) |
|---|---|---|
| Enzyme Kcat | 13,470 / 1,684 / 1,684 | 6,074 / 726 |
| Metal-ion binding | 6,000 / - / 1,332 | 3,263 / 660 |
| Thermostability (Human-only) | 1,979 / 247 / 248 | 1,818 / 202 |
| Thermostability (All) | 5,968 / 746 / 746 | 4,971 / 553 |
| Thermostability (Nanobody) | 522 / 95 / 147 | - |

**S. Table 2: Downstream benchmark dataset sizes.** Canonical columns give the original splits as obtained, including the validation set (HuggingFace parquet for enzyme Kcat & metal binding; for nanobody; the coverage-only single-domain Meltome split for thermostability). Single-domain columns give the seq-id 0.8 single-domain subset (MMseqs2 -c 0.8 --min-seq-id 0.8 vs TED365) used in the KNN benchmark: since a held-out validation set is meaningless for non-parametric k-NN zero-shot scoring, the original train and validation are merged into one training set, so only train/test are reported. Metal binding has no validation split. Nanobody was not single-domain filtered.

#### Jaccard-embedding distance analysis.

To provide a complementary, threshold-free view of how embedding similarity relates to functional similarity, we analysed the relationship between pairwise embedding distance and GO term overlap using the Jaccard Index on 100,000 randomly sampled non-homologous protein pairs from Swiss-Prot-GOA (2017). Result can be seen in Supplementary Figure 2. An AUROC analysis is also available in Supplementary Table 2.

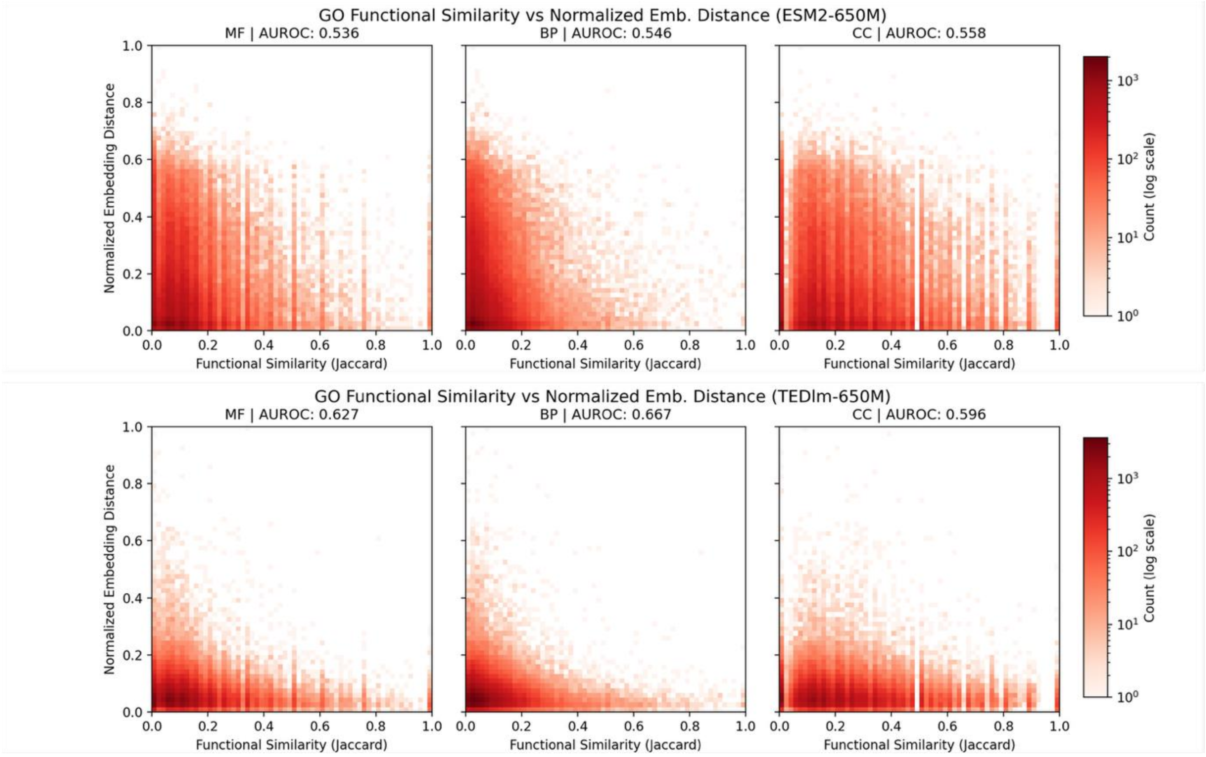


***S.Figure 7.*** *Function similarity (Jaccard) versus normalized embedding distance across Molecular Function (MF), Biological Process (BP), and Cellular Component (CC) namespaces. Each point represents a pair of proteins randomly sampled (100,000 pairs) from the SwissProt–GOA (2017) dataset, with functional similarity computed as the Jaccard Index of their GO term sets. Embedding distance defined as 1-norm(cos sim). The top row shows results for ESM2 650M, and the bottom for TEDlm 650M.*

**S. Table 3: Jaccard distance v.s. Embedding similarity. Performance** of ESM2 and TEDlm reported in AUROC across the MF/BP/CC namespaces. Data sampled randomly from CAFA3 set, in attempt to quantify the Jaccard distance GO semantic similarity plot in S.Figure 2.

| GO Namespace | ESM2 650M | ESM2 3B | TEDlm 650M | TEDlm 1.3B |
|---|---|---|---|---|
| MF AUROC | 0.536 | 0.519 | 0.627 | 0.616 |
| BP AUROC | 0.546 | 0.494 | 0.667 | 0.667 |
| CC AUROC | 0.558 | 0.506 | 0.596 | 0.584 |

1. Su, J. *et al.* RoFormer: Enhanced Transformer with Rotary Position Embedding. *Proceedings of the 2021 Conference on Empirical Methods in Natural Language Processing (EMNLP)* 7381–7392 (2021).

2. Dao, T. Flashattention-2: Faster attention with better parallelism and work partitioning. in *International Conference on Learning Representations* vol. 2024 35549–35562 (2024).

3. Shazeer, N., Parmar, N., Le, Q. & et al. GLU Variants Improve Transformer. *arXiv preprint arXiv:2202.10077* (2022).

4. Kendall, A., Gal, Y. & Cipolla, R. Multi-task learning using uncertainty to weigh losses for scene geometry and semantics. in *Proceedings of the IEEE conference on computer vision and pattern recognition* 7482–7491 (2018).

5. Loshchilov, I. & Hutter, F. Decoupled weight decay regularization. *arXiv preprint arXiv:1711.05101* (2017).

6. Paszke, A. *et al.* Pytorch: An imperative style, high-performance deep learning library. *Adv. Neural Inf. Process. Syst.* **32**, (2019).

7. Lin, Z. *et al.* Evolutionary-scale prediction of atomic-level protein structure with a language model. *Science (1979).* **379**, 1123–1130 (2023).

8. Senior, A. W. *et al.* Improved protein structure prediction using potentials from deep learning. *Nature* **577**, 706–710 (2020).
